# Insights into *Phakopsora pachyrhizi* effector-effector interactions

**DOI:** 10.1101/2023.08.30.555440

**Authors:** Mingsheng Qi, Haiyue Yu, Melissa Bredow, Aline Sartor Chicowski, Letícia Dias Fields, Steven A. Whitham

## Abstract

The multifaceted role of pathogen-encoded effectors in plant-pathogen interactions is complex and not fully understood. Effectors operate within intricate host environments, interacting with host proteins and other effectors to modulate virulence. The complex interplay between effectors raises the concept of metaeffectors, where some effectors regulate the activity of others. While previous research has demonstrated the importance of effector repertoires in pathogen virulence, only a limited number of studies have investigated the interactions between these effectors. This study explores the interactions among *Phakopsora pachyrhizi* effector candidates (*Pp*ECs). *P. pachyrhizi* haustorial transcriptome analysis identified a collection of predicted *Pp*ECs. Among these, *Pp*EC23 was found to interact with *Pp*EC48, prompting further exploration into their potential interaction with other effectors. Here, we utilized a yeast-two-hybrid screen to explore protein-protein interactions between *Pp*ECs. A split-luciferase complementation assay also demonstrated that these interactions could occur within soybean cells. Interestingly, *Pp*EC48 displayed the ability to interact with several small cysteine-rich proteins (SCRP), suggesting its affinity for this specific class of effectors. We show that these interactions involve a histidine-rich domain within *Pp*EC48, emphasizing the significance of structural motifs in mediating effector interactions. The unique nature of *Pp*EC48, showing no sequence matches in other organisms, suggests its relatively recent evolution and potential orphan gene status. Our work reveals insights into the intricate network of interactions among *P. pachyrhizi* effector-effector interactions.

## Main Text

Pathogen-encoded effectors promote susceptibility and also elicit defenses if their presence is detected by immune receptors (Yuan et al. 2021). Effectors function in complex settings within hosts in concert with dozens to potentially hundreds of other effectors (Kamoun 2006; Lanver et al. 2017; Lindeberg, Cunnac, and Collmer 2012; Lo Presti et al. 2015; Martin and Kamoun 2011; Petre, Joly, and Duplessis 2014; Stergiopoulos and de Wit 2009). The simultaneous presence of multiple effectors in host cells raises the possibility that direct and/or indirect effector-effector interactions are needed to bring about necessary changes in the host that promote virulence. Effectors may function redundantly with, cooperate with, assist, function as decoys for, or control other effectors to modulate host immunity. The identification of bacterial effectors of animal and plant pathogens that promote or antagonize the activity of other effectors has revealed the importance of effector interplay and given rise to the concept of metaeffectors (effectors of effectors) (Shames and Finlay 2012; Urbanus et al. 2016; Martel et al. 2022).

Given that filamentous eukaryotic pathogens of plants are predicted to encode many more effectors than bacterial pathogens (Lo Presti et al. 2017, 2015; Petre, Joly, and Duplessis 2014; Kamoun 2006), it is expected that there are also cooperative and hierarchical functional relationships among their effector repertoires. This is well exemplified by the *Phytophthora sojae* effector xyloglucan-specific endoglucanase (XEG1)-like protein 1 (*Ps*XLP1), which serves as an apoplastic decoy to bind a soybean glucanase inhibitor protein (*Gm*GIP1), which frees its paralog, *Ps*XEG1, to carry out its virulence functions in hydrolyzing cell wall polysaccharides (Ma et al. 2017). Various studies show that effectors may interact in a variety of ways and, in some cases, through physical interactions. For example, a large-scale yeast-two-hybrid (Y2H) screen found that 126 effector candidates from *Ustilago maydis* interacted with themselves or other effector candidates, representing almost a third of all effector candidates tested (Alcântara et al. 2019). Despite the overwhelming evidence that effector repertoires are important determinants of pathogen virulence and host range, only a few studies have systematically characterized the interactions that occur between these effectors.

Soybean rust (SBR) is a foliar disease caused by the obligate biotrophic fungus *Phakopsora pachyrhizi* (Bromfield 1984; Godoy et al. 2016). SBR occurs in all major soybean-growing countries and can cause yield losses of 80% or more (Kelly et al. 2015; Wrather et al. 2010). Despite the economic importance of *P. pachyrhizi* and other rust fungi, we lack a detailed understanding of the molecular mechanisms leading to disease. Rust fungi are obligate biotrophic basidiomycetes that must balance activities that promote fungal growth and development while keeping host cells alive and their defenses at bay (Voegele and Mendgen 2003). It seems logical that this would require finely-tuned regulation of effector activities that involve the cooperation and coordinated activity of the effectome.

Previously, sequencing of the haustorial transcriptome identified 156 haustoria-expressed secreted proteins, termed *P. pachyrhizi* effector candidates (*Pp*ECs) (Link et al. 2014). The open reading frames (ORFs) of 82 *Pp*ECs were cloned and expressed in heterologous systems to identify individual *Pp*ECs with effector-like functions. Seventeen *Pp*ECs suppressed pathogen-associated molecular pattern (PAMP)-triggered immunity (PTI) in *Nicotiana benthamiana* and Arabidopsis, including one (*Pp*EC23) that also suppresses bacterial-induced effector-triggered immunity (ETI) in *N. benthamiana, N. tabacum*, and soybean (Qi et al. 2016, 2018). We were interested in potential interactions that may occur among *Pp*EC23 and the other *Pp*ECs, which could influence their activities during infection. To test for protein-protein interactions occurring among these 82 *Pp*ECs, the open reading frames, minus the predicted signal peptides, were mobilized into the Y2H pGADT7 prey and pGBKT7-GW bait vectors and transformed into Y187 and AH109 yeast strains, respectively. The 82 AH109 and 82 Y187 clones were mated in order to test all possible combinations of interactions, including self-interactions, totaling 6,724 possible interactions.

To eliminate *Pp*ECs that autoactivate in the MatchMaker system (Takara Bio, Shiga, Japan), we tested each *Pp*EC in pGBKT7 or pGADT7 against the corresponding prey or bait plasmid, respectively, that was either empty or expressing the SV40 T-antigen for an additional 328 yeast matings. We observed growth of yeast carrying pGADT7-*Pp*EC54, -*Pp*EC63, -*Pp*EC21, -*Pp*EC47, -*Pp*EC51, and -*Pp*EC80 when mated with yeast transformed with the empty pGBKT7-GW or pGBKT7-T-antigen (Supplementary Fig. S1) suggesting that expression of the Gal4 promoter was auto-activated by these six *Pp*ECs. However, this auto-activation was not observed when the pGBKT7 vector was used to express these *Pp*ECs in yeast that was also carrying the pGADT7 empty or pGADT7-T antigen. In addition, we did not observe autoactivation of yeast growth by any other *Pp*EC expressed from pGBKT7. Based on these results, growth of yeast strains carrying pGADT7 expressing any of the six *Pp*ECs listed above was removed from consideration as indicating potential *Pp*EC*-Pp*EC interactions.

On triple dropout (TDO) media, pGADT7-*Pp*EC23/pGBKT7-*Pp*EC23 grew at both 2 and 4 days after plating (Supplementary Fig. S2), indicating that *Pp*EC23 forms at least dimers in yeast. Although evidence of the self-interaction was not observed on quadruple dropout (QDO) media (Supplementary Fig. S2), the results on TDO are consistent with our previous work showing that *Pp*EC23 self-interaction is mediated through its carboxy (C)-terminus (Qi et al. 2016). The pGADT7-*Pp*EC23 paired with pGBKT7-*Pp*EC63, -*Pp*EC65, -*Pp*EC75, and -*Pp*EC85 exhibited strong growth on TDO but not QDO. pGADT7-*Pp*EC23 paired with pGBKT7-*Pp*EC48 grew on both TDO and QDO (Supplementary Fig. S2), but the interaction was not observed when *Pp*EC23 and *Pp*EC48 were expressed from the bait and prey plasmids, respectively, indicating that the interaction is sensitive to the protein tag. This sensitivity is consistent with our previous observations that *Pp*EC23-*Gm*SPL12 complexes were strongly influenced by tag and orientation (Qi et al. 2016).

*Pp*EC48 interacted with six other *Pp*ECs in this screen: *Pp*EC59, *Pp*EC65, *Pp*EC75, *Pp*EC78, and *Pp*EC85 when expressed from either the bait or prey construct, but the interaction with *Pp*EC63 was only observed in the pGADT7-*Pp*EC63/pGBKT7-*Pp*EC48 configuration (Supplementary Fig. S2). Interestingly, we noted that these effector candidates, including *Pp*EC23, were the most cysteine-rich among this set of 82 candidate effectors, with a range of 6.59 to 8.33% cysteine (Supplementary Table S1). These effector candidates are also small, ranging from 116 to 291 amino acids, including the predicted signal peptides. *Pp*EC48 is not cysteine-rich and encodes two cysteine residues, one in the signal peptide and the other in the predicted mature protein. There was no evidence of *Pp*EC48 interacting with itself in the Y2H, which is consistent with its apparent affinity for small cysteine-rich proteins (SCRP). To investigate if effector-effector interactions identified by Y2H also occur in soybean cells, a split-luciferase complementation assay (SLCA) was used. The 11 *Pp*ECs of interest were mobilized by Gateway recombination (Invitrogen) into the pDuEx system in which fusions to the N and C terminal fragments of Renilla luciferase were created (Fujikawa and Kato 2007; Kato and Jones 2010). The constructs were transfected into soybean protoplasts, and after overnight incubation, the luciferase assay was performed. The SLCA positive control of Arabidopsis Histone 2A vs. Histone 2B produced the highest relative light units of any pair, indicating a strong interaction as expected (Fujikawa and Kato 2007). The negative control of *Pp*EC48 vs. GFP produced light levels similar to background, indicating non-detectable interaction. Most other pairwise combinations of *Pp*ECs tested produced relative light units that were significantly above background (Fig. 1A and Supplementary Fig. S3). The interactions that were identified only by Y2H or by a combination of Y2H and SLCA are summarized in Figure 1B. These data demonstrate that most of the effector-effector interactions observed in Y2H occurred in soybean cells at levels sufficient to be detected by SLCA, and they support the idea that *Pp*EC48 interacts with several SCRP.

**Figure 1.**
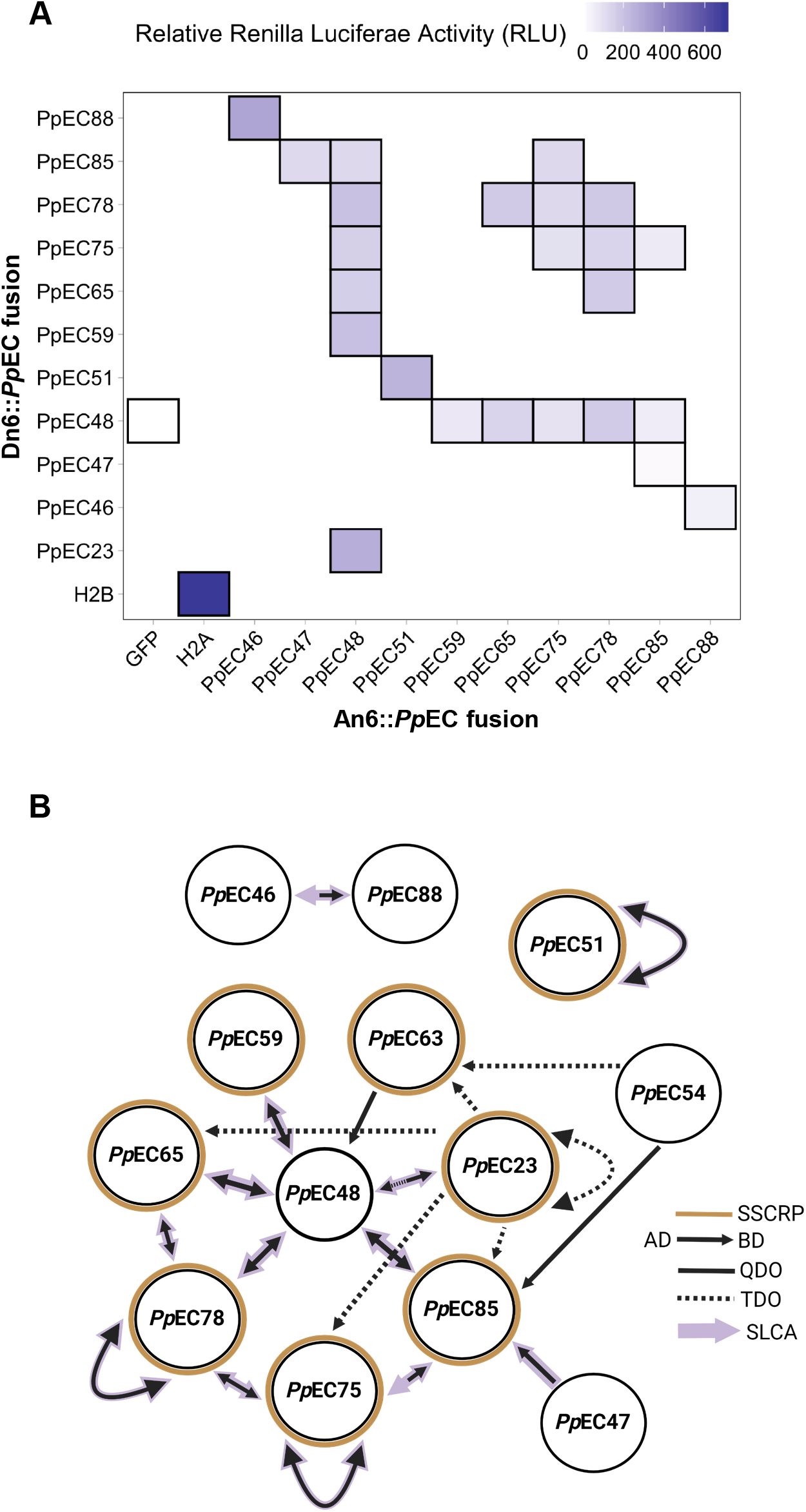
*Phakopsora pachyrhizi* effector-effector interactions network. **A**. Split-luciferase complementation assay to test interactions between *Pp*ECs in soybean protoplasts. Relative Renilla luciferase activity (RLU) was used to demonstrate interactions between *Pp*ECs as a result of their association and reconstitution of the two fragments of Renilla luciferase fused to each *Pp*EC in An6 or Dn6 vectors and breakdown of the ViviRen Live Cell substrate. Histone 2A (H2A) and histone 2B (H2B) were used as a positive control interaction, and green fluorescent protein (GFP) was used as a negative interaction control. **B**. Representation of the *Pp*EC-*Pp*EC interactions observed using the yeast-two-hybrid and split-luciferase complementation assays. SSCRP: Small secreted cysteine-rich proteins. AD: Activation domain; BD, DNA-binding domain; TDO: Triple drop-out medium (SD/–His/–Leu/–Trp); QDO: Quadruple drop-out medium (SD/– Ade/–His/–Leu/–Trp); SLCA: Split-luciferase complementation assay.

These results demonstrate that *Pp*EC48 has the ability to interact with several SCRPs in yeast. The temporal expression and subcellular localization of effectors is another important consideration that can help to understand effector-effector interactions. The *Pp*ECs of interest were all identified from mRNA transcripts that were present in haustoria extracted from infected soybean leaves at 10-14 days post-inoculation. This observation suggests that they are co-expressed, although it may be interesting to conduct detailed expression time course analyses in the future. We previously demonstrated that *Pp*EC48 and five of these interactors (*Pp*EC59, *Pp*EC63, *Pp*EC65, *Pp*EC78, and *Pp*EC85) localize to the nucleus when transiently expressed in *N. benthamiana* (Qi et al. 2018). *The co-expression in haustoria and detection of GFP-tagged proteins in the same compartment supports the hypothesis that Pp*EC48 can interact with these SCRPs in soybean cells during infection.

The predicted *Pp*EC48 protein is 204 amino acids, and analysis of the sequence using SignalP 5.0 (Almagro Armenteros et al. 2019) identifies a potential cleavage site between positions 24 and 25 with a probability of 0.8680. The predicted mature protein was subdivided into two domains, with domain A having no distinguishing features, while domain B is histidine-rich. BLASTp searches using the mature *Pp*EC48 amino acid sequence against Genbank nr return a sequence annotated as CSEP-04 from *P. pachyrhizi* (Kunjeti et al. 2016) *and PpacPPUFV02*|*3046184 (99*.*3% identity) from the recently sequenced UFV02 genome (Gupta et al. 2023). A comparison of the amino acid sequences of CSEP-04 to Pp*EC48 identifies a single amino acid polymorphism at position 26, suggesting that these proteins, which were identified in different *P. pachyrhizi* isolates, are encoded by the same gene. The lack of significant matches in any other organism, including other rust fungi for which genome sequence is available, suggests that *Pp*EC48 is unique to *P. pachyrhizi*. This species specificity indicates that *Pp*EC48 may be an orphan gene that has evolved relatively recently.

To identify the domain mediating interactions between *Pp*EC48 and the SCRP effector candidates, domains A and B were cloned into both the pGBKT7 (bait) and pGADT7 (prey) Y2H plasmids (Fig. 2A). These constructs were transformed into yeast, and the resulting strains were then mated with yeast strains carrying corresponding prey or bait plasmids expressing *Pp*EC23, *Pp*EC59, *Pp*EC65, *Pp*EC75, *Pp*EC78, and *Pp*EC85. Domain B interacted with all the SCRP in both the bait and prey configurations, and domain A only interacted with *Pp*EC78 in the pGADT7-*Pp*EC48A vs. pGBKT7-*Pp*EC78 configuration (Fig. 2B-C). These data indicate that the histidine-rich domain B is the primary mediator of the interactions of *Pp*EC48 with these six SCRP. Histidine-rich proteins have been identified in bacteria and fungi and have been associated with heavy metal binding, oxidative stress resistance, and macromolecule binding (Cheng et al. 2013; Nostadt et al. 2020). Proteins containing histidine-rich domains have also been associated with metal-dependent protein-protein interactions in plants (Heise et al. 2007), animals (Arvanitis et al. 2007), and apicomplexan parasites (Waller et al. 1999). We were therefore interested in identifying the interaction interfaces between *Pp*EC48B and the other interactors; however, we were unable to obtain a high-confidence model for *Pp*EC48 using AlphaFold2 (< 50% confidence) (Jumper et al. 2021), RoseTTAFold (52% confidence) (Advisory Board 2021), or Phyre 2.0 (48% confidence) (Kelley et al. 2015) algorithms, in line with its predicted classification as an orphan protein (Singh 2023).

**Figure 2.**
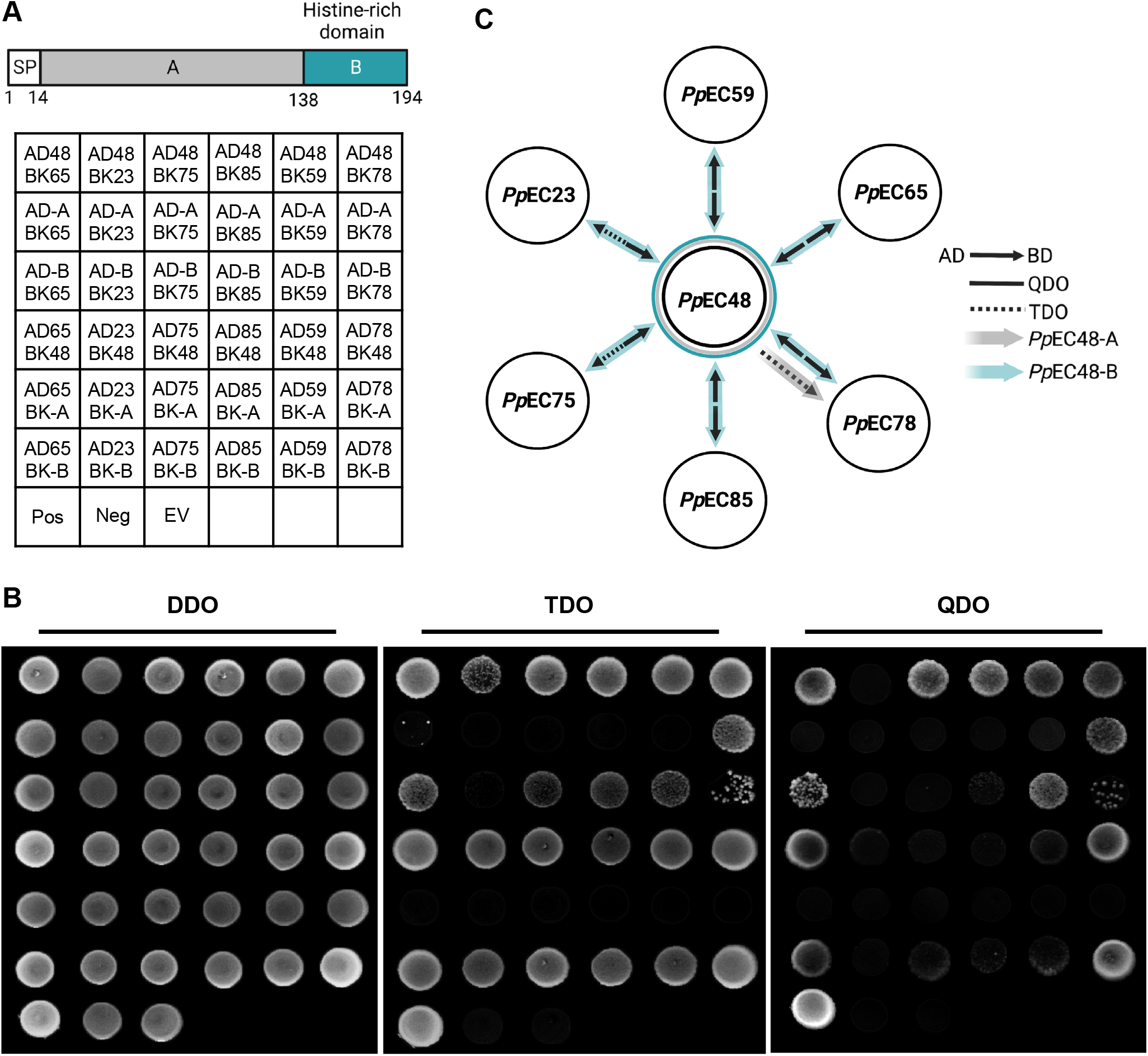
*Phakopsora pachyrhizi* effector candidate 48 (*Pp*EC48) functional domain analysis. **A**. Representation of *Pp*EC48 domains and yeast-two-hybrid plate organization. SP: signal peptide. AD: Activation domain. BK: DNA-binding domain. **B**. Yeast-two-hybrid screen demonstrating the interactions of *Pp*EC48 domains A or B with other *Pp*ECs. DDO: Double drop-out medium (SD/–Leu/–Trp); TDO: Triple drop-out medium (SD/–His/–Leu/–Trp); QDO: Quadruple drop-out medium (SD/–Ade/–His/–Leu/–Trp). **C**. Representation of the interactions between *Pp*EC48 domains A or B with other *Pp*ECs. AD: Activation domain; BD, DNA-binding domain; DDO: Double drop-out medium (SD/–Leu/–Trp); TDO: Triple drop-out medium (SD/– His/–Leu/–Trp); QDO: Quadruple drop-out medium (SD/–Ade/–His/–Leu/–Trp).

In this study, our overall goal was to develop a data set that moves us beyond studying single effectors in isolation to understanding roles for effector-effector complexes in *P. pachyrhizi* virulence. We identified several novel interactions between *Pp*ECs that could be of interest for functional analyses. One of these proteins, *Pp*EC48, interacts with a number of SCRPs, including *Pp*EC23, which suppresses basal defense responses in host and non-host species (Qi et al. 2016) (Supplementary Table S1). While *Pp*EC48 itself did not exhibit direct defense suppression activity in previous assays (Qi et al. 2018), its interactions with other effectors could indicate a regulatory role or participation in modulating host defenses. The ability of *Pp*EC48 to interact with several SCRPs suggests that it may function as a metaeffector. Future investigations could explore the functional implications of these interactions, their expression patterns, and how they collectively impact the virulence strategies of *P. pachyrhizi* during soybean rust infection. Using the EffectorP program to predict effector candidate proteins in *P. pachyrhizi*, Gupta et al. (2023) identified an average of 850 effector candidates in the genomes of the three isolates that were sequenced. Therefore, our study with 82 effectors covers approximately 1% of the total potential interactions, and it indicates that additional work to establish the *P. pachyrhizi* metaeffector repertoire will be interesting and necessary to fully understand the functions of effectors in this pathosystem. Overall, this study provides a stepping stone toward understanding the complex molecular mechanisms underlying plant-pathogen interactions and the interplay among effectors within the intricate host-pathogen landscape.

## Author contributions

M.Q., H.Y., S.A.W. conceived and planned the experiments; M.Q., H.Y., L.P.F. performed the experiments, collected and analyzed the data; M.B., A.S.C., M.Q., H.Y., S.A.W. interpreted the data; M.B., A.S.C., S.A.W. wrote the manuscript.

## Acknowledgments

We are grateful to Naohiro Kato (Louisiana State University) for providing the plasmids and protocols for the pDuEx split luciferase complementation assay, and to Yoshie Hanzawa (California State University, Northridge) for sharing protocols for soybean protoplasts. This work was supported in part by the Iowa State University Plant Sciences Institute, the Iowa Soybean Association, National Science Foundation Integrative Organismal Systems Plant Biotic Interactions program (IOS-1551452), USDA NIFA Hatch Project 4308, as well as fellowships from the China Scholarship Council (201706350083 to H.Y.), and Coordination for the Improvement of Higher Education Personnel (CAPES), Brazil to L.D.F.

## Supplementary Materials and Methods

### Bacterial and yeast strains and plasmids

Bacterial, yeast strains and plasmids used in this study are listed in Supplementary Table S1. *E. coli* and *Agrobacterium tumefaciens* were grown in Luria-Bertani (LB) broth at 37 °C (*E. coli*) or 28 °C (*A. tumefaciens*) using either liquid or solid media with appropriate antibiotics. *Saccharomyces cerevisiae* strains were grown at 30 °C in yeast extract peptone dextrose (YPD) medium or yeast minimal media with appropriate Dropout (DO) supplements using either liquid or solid media. Plasmids were introduced into *E. Coli, A. tumefaciens* and *S. cerevisiae* using heat shock, freeze-thaw and LiAc-PEG-mediated transformation methods (Gietz and Woods 2006), respectively.

### Plant growth conditions

Soybean (*Glycine max* cv. Williams 82) plants were grown in plastic pots in a controlled growth chamber under long-day conditions (16h of light and 8h of dark) at an average temperature of 23 °C, 400 μmol m^-2^s^-1^ irradiance, and 45 to 65% relative humidity.

### Manual array-based yeast two hybrid assays

Manual array-based yeast library construction and two-hybrid assay followed (Roberts et al. 2012) with minor modifications. Briefly, the 82 *Pp*EC ORFs without signal peptides were moved into pGBKT7-GW and pGADT7-GW through Gateway LR reaction (Invitrogen) and introduced into *E. coli*. Positive clones were confirmed with PCR and sequencing. The confirmed BD-*Pp*EC and AD-*Pp*EC plasmids were transformed into yeast AH109 and Y187 strains, respectively. The exhaustive one-to-one *Pp*EC interactions were screened using the normal Y2H approaches according to the manufacturer’s instructions (Clontech) except that most steps were adapted to 96-well plates. All resulting diploid clones were grown in liquid medium with selection for the two plasmids, and then cells were plated onto double drop out (DDO), triple drop out (TDO), and quadruple dropout (QDO) media. The DDO medium only selects for the presence of the bait and prey plasmids, and all clones grew demonstrating that transformation and subsequent mating was successful (e.g. Figure 1 DDO). The TDO selects for interactions due to complementation of histidine auxotropy and the more stringent QDO selects for complementation of histidine and leucine auxotrophy.

### *Pp*EC48 mutant derivatives

*Pp*EC48 amino acid sequence without signal peptide was manually examined via the Simple Modular Architecture Research Tool (SMART; http://smart.embl-heidelberg.de/) to support the idea that it is comprised of an unremarkable domain A and the histidine-rich domain domain B in addition to the N-terminal signal peptide. *Pp*EC48-A and *Pp*EC48-B were cloned using the primers listed in Supplementary Table S3.

### Split luciferase complementation assay in soybean protoplasts

Protoplasts were prepared as previously described with minor modifications (Wu and Hanzawa 2018). Newly expanded unifoliate leaves of 12-day-old soybean were cut into 0.5-1 mm strips using a new razor blade. The strips were gently transferred to a Petri dish with 10 ml of digestion buffer, containing 2% (w/v) Cellulase “onozuka” R-10 (Yakult), 0.1% Pectolyase Y-23 (Kyowa), 0.4 M mannitol, 20 mM MES (pH 5.7), 20 mM KCl, 10 mM CaCl_2_, 0.1% BSA and 0.5 mM DTT. The dish was then vacuum-sealed for 30 min at room temperature and incubated in the digestion buffer for 4 hrs in dark at room temperature with a gentle shake (speed: 30, tilt: 1) on a three-dimensional Rotator Waver (VWR International). As protoplasts were released, 10 ml of W5 buffer, containing 154 mM NaCl, 125 mM CaCl_2_, 5 mM KCl, 2 mM MES (pH 5.7), was added to the Petri dish. The enzyme/protoplast solution was filtered with 75-μM nylon mesh and transferred to a 50 ml round-bottom tube. The protoplasts were collected by centrifuging at 100 x g for 3 min. The protoplast pellets were washed once in 10 ml of W5 buffer followed by centrifuging at 100 x g for 3 min. Then the protoplasts were resuspended in 10 ml of W5 solution and kept on ice for 30 min. After centrifuging and removal of the supernatant, the protoplasts were resuspended with an appropriate volume of MMG solution, containing 0.4 M mannitol, 4 mM MES (pH 5.7), 15 mM MgCl_2_, to the final concentration at 10^6^ cells/ml by counting with a hemocytometer. 50 μg of plasmid DNA and 500 μl of protoplast suspension (5 × 10^5^ cells) were dispensed into each 15 ml tube. Then, 550 μl of PEG solution, containing 40% (w/v) PEG4000, 0.2 M mannitol, 100 mM CaCl_2,_ was added and gently shaken for 30 sec. After incubating the protoplast cells for 15 min at room temperature, 2 ml of W5 buffer was added to stop the transfection. After centrifuging and removal of the supernatant, 500 μl of WI solution, containing 0.5 M mannitol, 4 mM MES (pH 5.7), 20 mM KCl, was added to the tube. Then the protoplast cells were resuspended and transferred into a 6-well tissue culture plate (Corning) for overnight incubation under low fluorescent light (4 μmol m^-2^ s^-1^) at room temperature. The transfected protoplast cells were harvested by low-speed centrifugation (100 x g for 3 min) and were resuspended in 50 μl of WI solution. For the luminescence detection, ViviRen Live Cell substrate (Promega), a coelenterazine derivative, was used as the substrate. The ViviRen stock solution (6 mM) was diluted 100 times with W5 buffer to form the working solution and 10 μl of ViviRen working solution was mixed with 50 μl of WI-resuspended protoplast cells for 30 min at room temperature in dark before measuring the luminescence. The luminescence signal of each sample was detected with a BioTek Synergy HT plate reader.

## Supplementary materials

### Supplementary Materials and Methods

**Figure S1.**
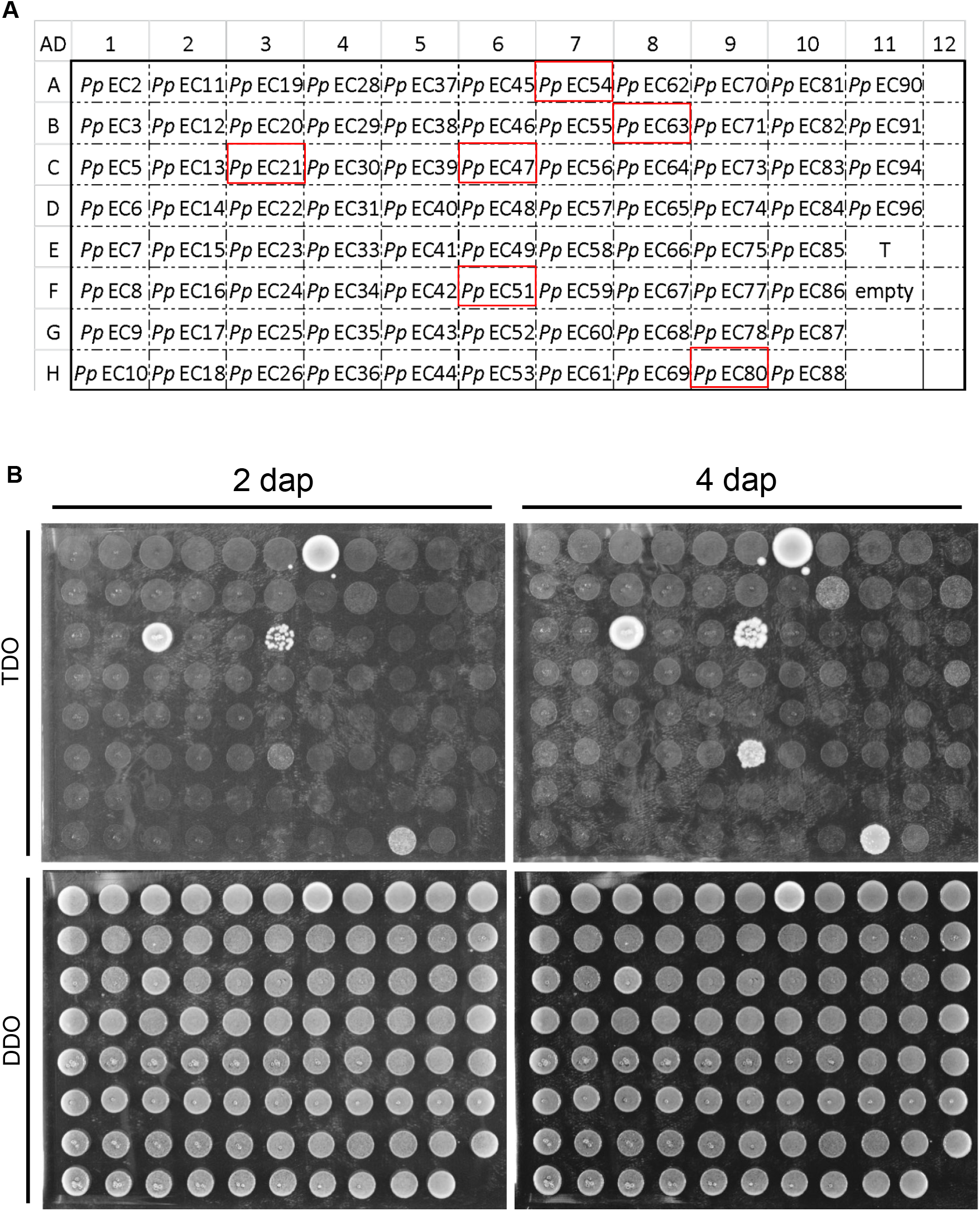
Yeast-two-hybrid autoactivation test of target *Pp*EC proteins. **A**. Plate organization and *Pp*ECs used to test for autoactivation of Y2H-inducible reporter genes. *Pp*ECs outlined in red were considered to autoactivate when fused to the activation domain, and their potential for interaction was excluded from consideration. **B**. Growth of yeast cells carrying the each *Pp*EC fused to the activation domain at 2 and 4 days after plating (dap). DDO: Double Drop-Out medium (SD/–Leu/–Trp); TDO: Triple Drop-Out medium (SD/–His/–Leu/–Trp).

**Figure S2.**
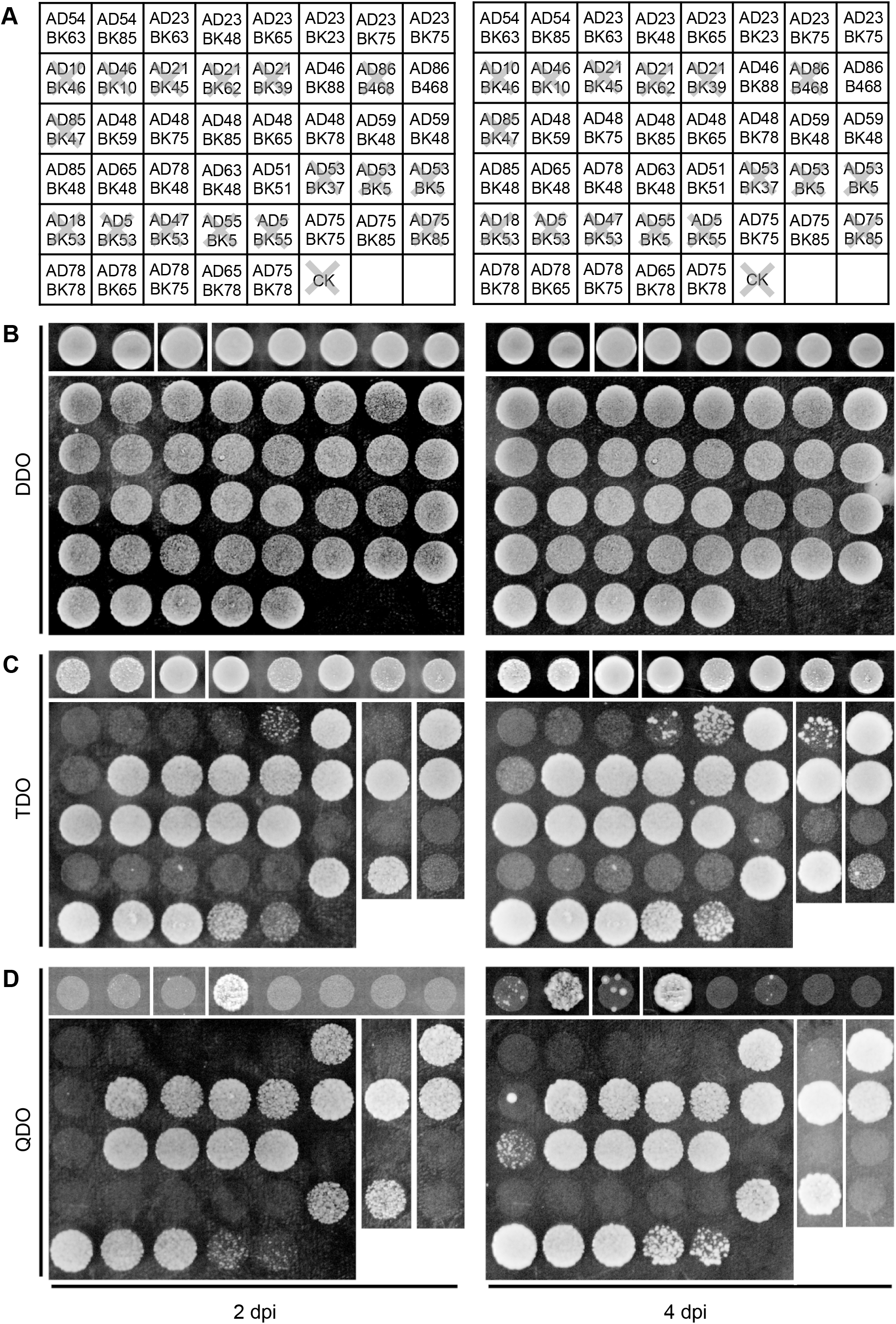
*Phakopsora pachyrhizi* effector-effector interactions identified by yeast-two-hybrid. **A**. Representation of the confirmed direct interactions between *Pp*ECs on the plates. AD: Activation domain; BK, DNA-binding domain. X indicates that results from selective media were interpreted as no growth. **B**. DDO: Double Drop-Out medium (SD/–Leu/–Trp); **C**. TDO: Triple Drop-Out medium (SD/–His/– Leu/–Trp); QDO: Quadruple Drop-Out medium (SD/–Ade/–His/–Leu/–Trp).

**Figure S3.**
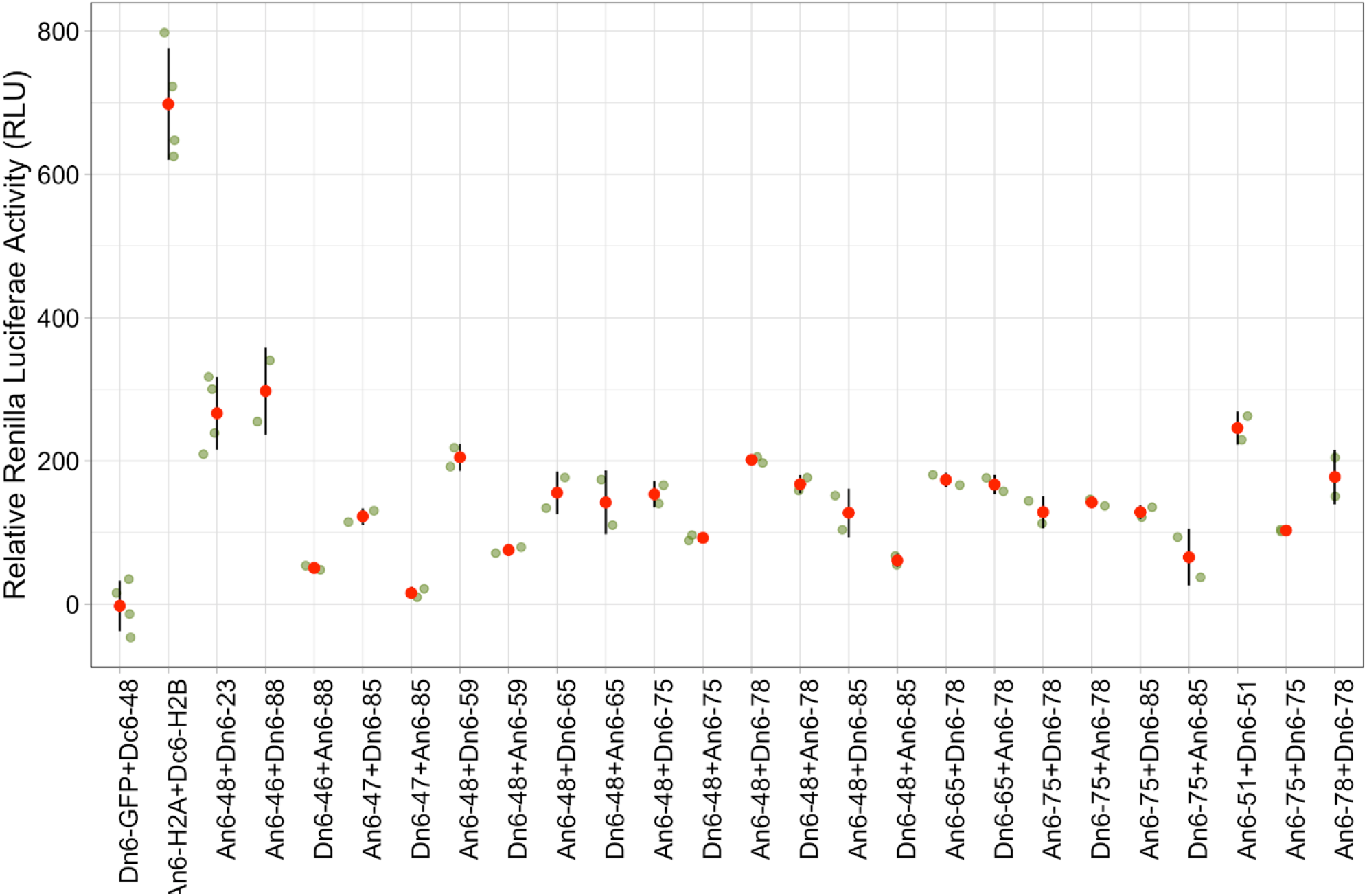
Split-luciferase complementation assay between *Pp*ECs. Relative Renilla luciferase activity (RLU) was used to demonstrate interactions between *Pp*ECs as a result of their association and reconstitution of the two fragments of Renilla luciferase fused to each *Pp*EC in pDuEx-An6 or pDuEx-Dn6 vectors, and breakdown of the ViviRen Live Cell substrate. H2A and H2B were used as a positive control pair, and GFP was used as a negative control. These data are depicted in heat map form in Figure 1A.

**Table S1.** Properties of *Pp*EC proteins identified in this study. Blue highlighted rows indicate the small cysteine-rich *Pp*ECs that interact with *Pp*EC48.

| *PpEC* No. | Annotation | Link et al. 2014 lineage specificity | Length (aa) | Cysteine No. | Cysteine% | Subcellular localization | Δbasal defense | Δcallose deposition | ΔHR | Yeast cell death suppression | Pepper HR induction | |
| --- | --- | --- | --- | --- | --- | --- | --- | --- | --- | --- | --- | --- |
|  |  |  |  |  |  |  |  |  |  |  | CM334 | ECW-30R |
| *PpEC23* | CSEP-35 (Kunjetti et al. 2016) | Pucciniales | 291 | 21 | 7.22 | c | +++ | +++ | + | Negative | na | na |
| *PpEC46* | hypothetical protein | Pucciniales | 207 | 2 | 0.97 | nc | - | - | - | Strong | - | - |
| *PpEC47* | hypothetical protein | no | 206 | 4 | 1.94 | n | - | - | - | Medium | - | na |
| *PpEC48* | CSEP-04 (Kunjetti et al. 2016) | Pucciniales | 192 | 2 | 0.98 | n | - | - | - | Medium | - | na |
| *PpEC51* | CSEP-08 (Kunjetti et al. 2016) | Pucciniales | 198 | 12 | 6.06 | n | - | - | - | Medium | - | - |
| *PpEC54* | No hits | Pp | 193 | 1 | 0.52 | nc | - | - | - | Weak | - | - |
| *PpEC59* | CSEP-09 (increase oomycetes virulence) | Pucciniales | 182 | 12 | 6.59 | n | - | - | - | Strong | - | - |
| *PpEC63* | hypothetical protein | Pucciniales | 168 | 14 | 8.33 | n | - | - | - | Strong | na | na |
| *PpEC65* | hesp-C49 (Kunjetti et al. 2016) | Pucciniales | 163 | 12 | 7.36 | n | - | - | - | Medium | - | + |
| *PpEC75* | hypothetical protein | Pucciniales | 133 | 11 | 8.27 | c | - | - | - | Strong | na | na |
| *PpEC78* | hypothetical protein | Pucciniales | 127 | 10 | 7.87 | n | - | - | - | Strong | - | - |
| *PpEC85* | hypothetical protein | Pucciniales | 116 | 8 | 6.9 | nc | - | - | - | Strong | - | + |
| *PpEC88* | hypothetical protein | no | 111 | 2 | 1.8 | n | - | - | - | Medium | na | na |

**Table S2.** Microbe strains and plasmids used in this study.

| Strain or plasmid | Genotype or relevant phenotype * | Source or reference |
| --- | --- | --- |
| *E. coli* |  |  |
| DH5α | F<sup>-</sup> *endA1 glnV44 thi-1 recA1 relA1 gyrA96 deoR nupG* Φ80d*lacZ*ΔM15 Δ(*lacZYA-argF*)U169 *hsdR17*(r<sub>K</sub><sup>-</sup> m<sub>K</sub><sup>+</sup>) λ<sup>-</sup> | Invitrogen |
| TOP10 | F<sup>-</sup> *mcrA* Δ(*mrr-hsdRMS-mcrBC*) Φ80*lacZ*ΔM15 Δ*lacX74 nupG recA1 araD139* Δ(*ara-leu*)7697 *galE15 galK16 rpsL*(Sp<sup>r</sup>) *endA1* λ<sup>-</sup> | Invitrogen |
| *Agrobacterium tumefaciens* |  |  |
| GV3101 | Carries Vir plasmid encoding T-DNA transfer machinery, Rif<sup>r</sup>, Gm<sup>r</sup> | Lab stock |
| *Saccharomyces cerevisiae* |  |  |
| AH109 | *MATα, trp1-901, leu2-3, 112, ura3-52, his3-200, gal4Δ, gal80Δ, LYS2::GAL1<sub>UAS</sub>-GAL1<sub>TATA</sub>-HIS3, GAL2<sub>UAS</sub>-GAL2<sub>TATA</sub>-ADE2, URA3::MEL1<sub>UAS</sub>-MEL1<sub>TATA</sub>-lacZ* | Clontech |
| Y187 | *MATα, ura3-52, his3-200, ade2-101, trp1-901, leu2-3, 112, gal4Δ, met<sup>-</sup>, gal80Δ, URA3::GAL1<sub>UAS</sub>-GAL1<sub>TATA</sub>-lacZ* | Clontech |
| Plasmids |  |  |
| pCR®8GW/TOPO® | Gateway-compatible entry vector, Sp<sup>r</sup> | Invitrogen |
| pGBKT7-GW | Gateway-compatible version of pGBKT7, Km<sup>r</sup> | Melissa G. Mitchum Lab |
| pGADT7-GW | Gateway-compatible version of pGADT7, Km<sup>r</sup> | Melissa G. Mitchum Lab |
| pDuExAn6-GW | Gateway recombination will result in a vector that express NLuc-Target Protein, Amp<sup>r</sup> | (Fujikawa and Kato et al.2007) |
| pDuExAc6-GW | Gateway recombination will result in a vector that express Target Protein-NLuc, Amp<sup>r</sup> |  |
| pDuExDn6-GW | Gateway recombination will result in a vector that express CLuc-Target Protein, Amp<sup>r</sup> |  |
| pDuExDc6-GW | Gateway recombination will result in a vector that express Target Protein-CLuc, Amp<sup>r</sup> |  |

**Table S3.**
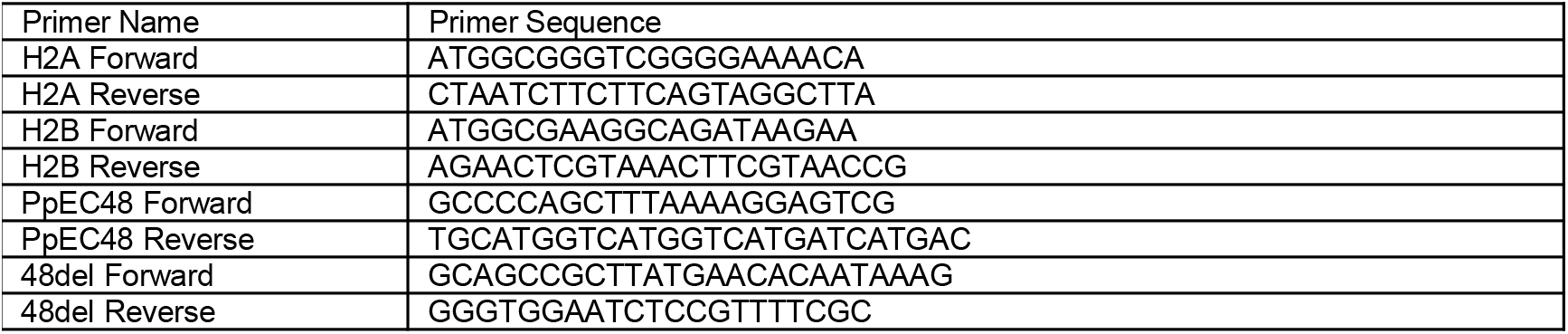
Oligonucleotide primers used in this study.

